# Autoregulation of the MET Receptor Tyrosine Kinase by its Intracellular Juxtamembrane Domain

**DOI:** 10.1101/2025.10.19.683305

**Authors:** Edmond M. Linossi, Carla A. Espinoza, Gabriella O. Estevam, James S. Fraser, Natalia Jura

**Affiliations:** Cardiovascular Research Institute, University of California San Francisco, San Francisco, CA 94158, USA; Department of Cellular and Molecular Pharmacology, University of California San Francisco, San Francisco, CA 94158, USA; Quantitative Biosciences Institute, University of California, San Francisco, San Francisco, CA 94158, USA; Department of Bioengineering and Therapeutic Sciences, University of California, San Francisco, San Francisco, CA 94158, USA

**Keywords:** MET receptor tyrosine kinase, HGF, hepatocyte growth factor, juxtamembrane domain, kinase regulation, phosphorylation, oncogenic mutations

## Abstract

Receptor Tyrosine Kinases (RTKs) are single-pass transmembrane receptors whose activation is tightly regulated by intra-domain interactions within both their extracellular and intracellular regions. The intracellular juxtamembrane domain, which links the transmembrane and kinase domains, often plays a critical role in modulating kinase activity. The MET receptor, activated by Hepatocyte Growth Factor (HGF), requires precise regulation to support normal development and wound healing, but becomes a potent oncogene when overexpressed or mutated. A common oncogenic lesion in MET, caused by exon 14 skipping, leads to partial deletion of its unusually long intracellular juxtamembrane domain and is frequently detected in non-small cell lung cancer (NSCLC), as well as pancreatic, liver and brain cancers. Despite its length and abundance of post-translational modifications, the functional role of the MET juxtamembrane domain has remained poorly understood. We have uncovered that this segment regulates the kinetics of MET kinase activation. Specifically, we found that a membrane-proximal, N-terminal region of the juxtamembrane domain accelerates activation loop phosphorylation promoting kinase transition to an active state. This regulation does not depend on the oligomeric state of MET but likely acts allosterically to enhance autophosphorylation of the kinase domain. Notably, this function is absent in the closely related MST1R/RON RTK, suggesting it is a unique feature of the MET receptor. Together, these findings uncover a previously unrecognized layer of MET regulation with potential implications for the development of selective therapies targeting MET-driven cancers.

## Introduction

Single-pass transmembrane Receptor Tyrosine Kinases (RTKs) lie at the apex of cellular signaling cascades. Binding of cognate growth factors to the extracellular domains induces RTK multimerization, or reorganization of existing receptor dimers, which in turn activates the intracellular tyrosine kinase domains (1). The activated kinase domains then proceed to phosphorylate tyrosine residues in receptor intracellular domains, which recruit signaling molecules that frequently also undergo phosphorylation, triggering a spectrum of downstream signaling cascades (2). Those include cellular motility, differentiation and proliferation pathways, among others. The MET receptor, an RTK activated by the hepatocyte growth factor (HGF), plays a central role in regulating cell motility and proliferation (3–5). MET signaling is essential for normal development in multicellular organisms and contributes to wound healing and tissue remodeling in adults (6, 7). A growing body of evidence also points to key roles of MET in immune and neuronal cell functions (8–10). Aberrant activation of MET via gene amplification or mutations is a prominent feature of multiple human cancers (11, 12).

MET contains a canonical tyrosine kinase domain, composed of an N-lobe and a C-lobe with the catalytic active site located in between (13). Multiple crystal structures of the MET kinase domain and *in vitro* enzymatic studies have outlined the key steps of MET kinase activation (14). In the inactive state, MET adopts a Src/CDK-like inactive conformation in which helix αC is swung away from the active site and the activation loop adopts a short helical fold blocking substrate access. Sequential phosphorylation of the activation loop tyrosines in MET (Y1234 and Y1235) is key to the release of this autoinhibition, leading to the extension of the activation loop and enabling substrate binding (13, 15–18). This step also allows rotation of helix αC towards the active site, establishing an essential Lys - Glu salt bridge (K1110 and E1127 in MET) that coordinates ATP for hydrolysis. In that way, MET follows a mechanism shared by other RTKs, including Insulin Receptor (IR), Platelet-Derived Growth Factor Receptor (PDGFR), FMS-like Tyrosine Kinase 3 (FLT3) and c-Kit, that rely on activation loop transphosphorylation for activity (19, 20).

Additional regulatory mechanisms that maintain MET kinase in its autoinhibited state have been identified, with many of them compromised by mutations in cancer. For example, residues Y1230 and D1228 in the activation loop, and C-lobe residues F1200 and L1195, help stabilize the inactive conformation of the activation loop and their mutation likely contribute to the disruption of the autoinhibitory tether (13, 17). Somewhat unexpectedly, these mutations often fail to induce constitutive receptor signaling and instead, they retain MET dependence on HGF binding for activation, even in an oncogenic setting (21–24). Characterization of these mutations *in vitro* using recombinant MET kinase domain revealed that they accelerate kinase activation without altering the maximal catalytic rate of the enzyme (25). These findings suggest that lowering the activation barrier for kinase activation alone is sufficient to drive oncogenic signaling, underscoring the importance of built in autoinhibitory mechanisms in preventing untimely MET signaling.

One of the most common MET oncogenic mutations, occurring in approximately 5% of non-small cell lung cancer (NSCLC), falls outside of the kinase domain and results in skipping of exon 14, which encodes the N-terminal half of the MET intracellular juxtamembrane (JM) domain (26–29). This disease mutation underscores potential importance of the JM domain, which connects the kinase domain to the transmembrane domain, in MET regulation. In many RTKs, JM domains serve as central regulatory elements controlling kinase activity (30). For example, in the inactive PDGFR, c-KIT, FLT3 and erythropoietin-producing human hepatocellular (Eph) receptors, the JM domain is directly tethered to the kinase domain restricting rotation of helix αC (31–34). This autoinhibitory mechanism is released by JM domain phosphorylation during activation of each of these receptors. In the human epidermal growth factors receptors (EGFR/HERs), the JM domain does not seem to play a critical inhibitory role, but it is essential for kinase dimerization and allosteric control of kinase activation upon growth factor stimulation (35, 36). In the Insulin Receptor (IR), which is a constitutive dimer, the JM domain switches from an inhibitory segment to an activating one upon IR stimulation (37–39). By comparison, the role of the JM domain in regulation of MET kinase activity remains poorly characterized.

In MET, the JM domain is substantially longer than in other RTKs: 92 residues in human MET (amino acids: 956-1048) compared to typical ∼10-50 amino acid length (30). While a direct role of the JM domain in regulating MET kinase activity has not been established, specific residues within this region have been shown to contribute to negative feedback regulation of MET receptor signaling. Phosphorylation of Y1003 results in recruitment of the c-Cbl E3 ligase followed by ubiquitination and lysosomal degradation of MET (40, 41). Phosphorylation of S985 by PKC also negatively regulates MET, although the underlying mechanism remains unknown (42, 43). Under cellular stress, the MET JM domain can be cleaved by caspases at D1000 to release an intracellular MET fragment that promotes pro-apoptotic signaling (44–46). Oncogenic exon 14 skipping mutations, which result in an in-frame deletion of residues 963-1009, eliminate all of these regulatory sites, complicating understanding how JM domain contributes to MET-driven oncogenesis (47). Elucidating the mechanisms by which the JM domain regulates MET kinase activity is therefore critical for understanding both its physiological and pathological roles.

In this study, we have investigated the direct role of the JM domain in regulation of MET kinase activity using purified proteins *in vitro*. Our findings reveal that the JM domain accelerates the autophosphorylation reaction required for full MET kinase activity, and that this activity maps to the membrane-proximal, N-terminal region of the JM domain that largely overlaps with exon 14-encoded sequence. In contrast to other RTKs, the activating effect of the JM domain is not exerted by promoting kinase oligomerization, but appears to entail allosteric regulation of the kinases domain. Moreover, this mechanism does not rely on the Y1003 or S985 sites, suggesting that the activating role of the JM domain is distinct from its previously known regulatory functions in MET signaling. We also demonstrate that the activity of the kinase domain of RON, a closely related MET ortholog, is not directly regulated by the JM domain, suggesting that MET employs a distinct, JM-driven mechanism for positive kinase regulation within the RTK family.

## Results

### The juxtamembrane region accelerates MET autophosphorylation

We investigated if the activity of the recombinant MET kinase domain *in vitro* is modulated by the presence of the JM domain (residues K956 - Q1048) (**Figure 1A**). We also looked if the presence of a short C-terminal tail (C-tail) in MET, which encompasses residues G1346-S1390 (**Figure 1A**), contributes to potential JM domain-dependent regulation. A series of recombinant MET intracellular constructs were generated to include or omit the JM domain or C-tail: the full intracellular domain including the C-tail (ICD: residues K956-S1390) or without the C-tail (ICD^ΔC-tail^: residues K956 - G1346), kinase domain with the C-tail (KD: residues Q1048-S1390) or without the C-tail (KD^ΔC-tail^: residues Q1048-G1346). All constructs were expressed in insect cells, purified and subject to dephosphorylation by lambda protein phosphatase to remove any tyrosine, serine or threonine phosphorylation that could potentially elevate basal kinase activity. Kinase activity of all constructs was monitored in the absence or presence of a generic poly (Glu:Tyr) 4:1 substrate (hereafter referred to as “substrate”) using a coupled kinase assay (35, 48) to evaluate autophosphorylation and substrate-directed MET activity.

**Figure 1.**
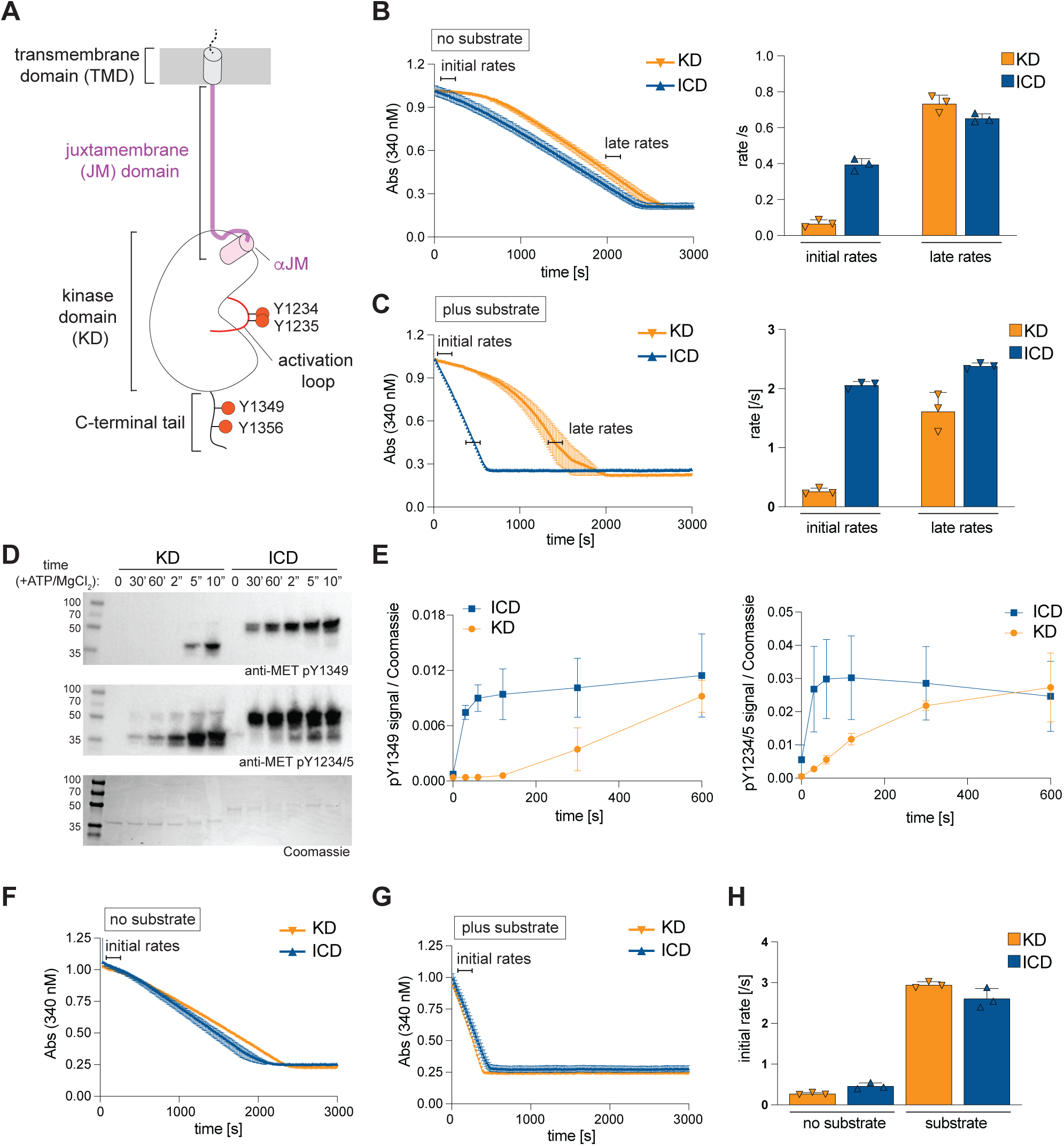
The intracellular juxtamembrane domain accelerates MET kinase domain activation. **(A)** Schematic representation of the MET receptor intracellular and transmembrane domains. The extracellular ligand binding region of MET is not shown. Regulatory phosphorylation sites in the activation loop and in the C-terminal tail are marked. (**B-C**) On the left, representative traces from the coupled kinase assay depict ATP consumption over time for the dephosphorylated kinase domain (KD) and full intracellular domain (ICD). The indicated regions used to calculate initial and late catalytic rates are marked by horizontal lines, and the rates are plotted in bar graphs on the right (triplicate data from one representative experiment). Measurements were conducted without the poly (Glu:Tyr) 4:1 substrate and correspond to KD autophosphorylation (B), or in the presence of a substrate (C). **(D)** Western blot analysis of KD and ICD autophosphorylation reaction using phospho-specific antibodies (specified below each blot). Coomassie staining was used to verify protein levels in all samples. **(E)** Quantification of the phosphorylated MET signal (Western) relative to total MET protein (Coomassie) for each site from three independent experiments. (**F-G**) Representative traces from the coupled kinase assay for the phosphorylated KD and ICD constructs, obtained in the absence (F) or presence of the substrate (G). (**H**) The calculated catalytic rates from triplicate measurements for representative experiments are shown in (F-G). Error bars represent standard deviation (SD). Supplementary Table 1 summarizes average values for each kinase assay from independent experiments.

The enzymatic activity of the KD displayed biphasic behavior during both autophosphorylation and substrate phosphorylation reactions, characterized by a slow initial rate followed by a faster rate, which we refer to as the “late” rate (**Figure 1B & C**). Although the ICD and KD late rates were comparable, we observed that the presence of the JM domain led to acceleration of both MET kinase autophosphorylation and substrate phosphorylation. We consistently measured a ∼4.5-fold faster autophosphorylation rate k_cat_ of the ICD compared to the KD (0.104 ± 0.023 s^-1^ compared to 0.466 ± 0.036 s^-1^, respectively) (**Figure 1B**). This difference in activity was further amplified (∼9-fold faster) in the presence of the substrate (KD k_cat_ = 0.168 ± 0.019 s^-1^ compared to 1.546 ± 0.172 s^-1^ for ICD) (**Figure 1C**). These results show that the JM domain accelerates the kinetics of activation without altering the enzyme’s maximal catalytic rate.

Biphasic activation of dephosphorylated KD was previously reported (49), with a transition between slow initial rate and a faster late rate attributed to autophosphorylation of the MET activation loop on tyrosines Y1234 and Y1235. To investigate if KD and ICD exhibit different kinetics of activation loop phosphorylation, we dephosphorylated these constructs followed by incubation with ATP and MgCl_2_ and monitored the phosphorylation of the activation loop tyrosines Y1234 and Y1235, as well as the C-tail tyrosine Y1235, over time by Western blot. Compared to the ICD, the KD exhibited a substantially slower rate of phosphorylation for both activation loop and C-tail sites (**Figure 1D & E**). These results show that the observed slower catalytic rate of the KD in the absence of the JM domain is due to delayed activation loop phosphorylation. If this is indeed the mechanism, we hypothesized that the KD construct in which the activation loop is fully phosphorylated prior to kinase activity measurements would not exhibit biphasic activation, and have a comparable catalytic rate to ICD. To test this, the ICD and KD constructs were pre-incubated with ATP and MgCl_2_ prior to the kinase assay. Under these conditions, KD activation curve was no longer biphasic, and the catalytic rates of KD and ICD were comparable in the presence or absence of the substrate (**Figure 1F - H**). These results further support our conclusion that the JM domain augments MET kinase autophosphorylation, accelerating the initial rate of its activation.

The presence of the C-tail did not change the existing differences between initial rates of the KD and ICD constructs: the rate of MET ICD^ΔC-tail^ was significantly faster compared to the MET KD^ΔC-tail^ for both autophosphorylation and substrate-dependent activity (**Supp. Figure 1A & B**). Removal of the C-tail from the ICD resulted in a small increase in catalytic rate relative to the construct containing the intact C-tail (1.7-fold increase), pointing to a minor but reproducible negative regulation of the full ICD by the C-tail. Previous studies reported enhanced MET receptor activation in cells upon C-tail removal (50, 51), and our findings with purified protein suggest this results from direct inhibition of the kinase domain by the C-tail. Nevertheless, the mechanisms by which JM domain and C-tail regulate MET kinase activity appear to be independent, as the presence of the JM domain accelerated initial rates under all conditions.

### *In silico* sequence and structural analysis of the MET JM

To investigate the mechanisms for the potentiating effect of the JM domain on kinase activation, we performed an *in silico* analysis of its sequence conservation and potential secondary structural elements. In these predictions we included the most C-terminal region of the JM domain (residues 1049-1070), which includes a short helix that is observed in all MET kinase domain crystal structures and is likely integral to the kinase domain fold (13). We recently denoted this region as helix αJM (52). The remaining MET JM domain encodes 92 amino acids (residues 956-1048) and has never been structurally characterized. For the purposes of the analysis, we divided this region into two segments: JM1 and JM2 (**Figure 2A**). The N-terminal segment (residues K956-D1010), designated as JM1, includes a short poly-basic sequence (residues K956-K962), predicted to bind the negatively charged lipids at the inner leaflet of the plasma membrane (53), and the region encoded by exon 14 (residues D963-E1009). The JM1 also includes two regulatory phosphorylation sites, a c-Cbl recruitment site (Y1003) (40) and the PKC substrate residue S985 (43). The C-terminal segment (residues D1010-Q1048) was designated as JM2 (**Figure 2A**).

**Figure 2.**
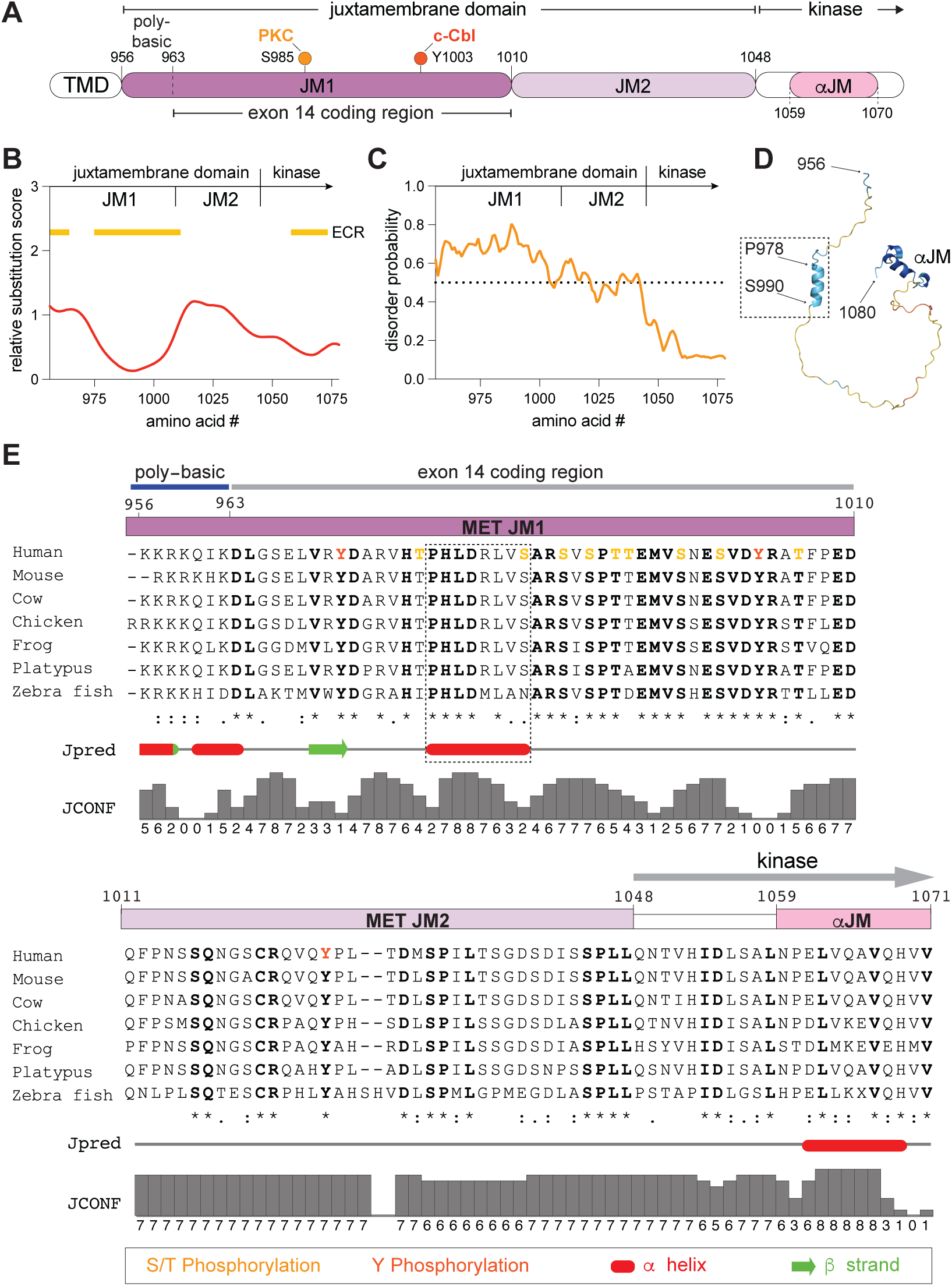
Evolutionarily conserved features of the MET juxtamembrane domain. **(A)** Linear domain schematic of the MET intracellular fragment with key regions and amino acid numbers annotated. Key regulatory phospho-sites are indicated for reference: S985, which is phosphorylated by PKC, and Y1003, which recruits c-Cbl when phosphorylated. TMD stands for the transmembrane domain **(B)** Conservation of the MET JM sequence assessed using the *Aminode* algorithm. Values on Y-axis represent relative substitution score for each amino acid number, with higher numbers indicating less sequence conservation. Yellow bars mark evolutionary conserved regions (ECRs) within the JM. **(C)** Disorder prediction for the MET JM using the IUPred2A algorithm. Dotted line indicates order-disorder threshold; higher values reflect greater disorder. N-terminal region of the kinase domain fold is included for comparison. **(D)** AlphaFold3 model of the MET JM domain, colored by pLDDT (predicted local distance difference test) confidence scores. The location of the predicted N-terminal JM helix, the helix αJM and boundary residues of the N- and C-terminal JM residues are annotated. **(E)** JM domain sequence alignment of selected MET orthologs is shown with subdomain regions annotated as in (A). Known phosphorylation sites are marked in red or orange. Dotted box marks the predicted N-terminal JM helix. Relative secondary structure predictions (marked red for α helix and green for β sheet) and the corresponding confidence scores are shown below the sequence alignment.

We assessed sequence conservation of the MET JM domain using Aminode (54), which determines evolutionary constrained regions (ECRs) that represent contiguous sections of higher amino acid sequence conservation with respect to phylogenetic distance between different MET orthologs (**Figure 2B**). This analysis showed that the JM domain contains two ECRs: the poly-basic region that directly follows the transmembrane helix and a portion of JM1 (residues V975-D1010), which includes most of the exon 14 coding region (**Figure 2B**). Conversely, the JM2 segment displayed lower evolutionary sequence conservation and a higher relative substitution score. Sequence alignment of MET orthologs also highlighted higher conservation of sequence and length across the JM1 segment compared to JM2 (**Figure 2E**; **Supp. Figure 2** displays sequence conservation across all orthologs). Additionally, phosphorylation sites within the JM domain, as annotated in PhosphoSitePlus database (55), are predominantly clustered in the JM1 region (**Figure 2E**). Although the entire JM domain is predicted to be largely disordered (**Figure 2C**), both the Jpred secondary structure prediction tool (56) and AlphaFold3 (57) predict an isolated, short alpha helix in the JM1 segment (JpreD: residues P978-S985; AlphaFold3: residues P978-S990) that has not been previously described (**Figure 2D & E**). Collectively, these analyses highlight that the JM1 segment of the MET JM domain is highly conserved, has evolved as a platform for post-translational modifications, and contains a region with a propensity to adopt a short helix. Together, these features point to JM1 as a region with the potential to act as regulatory control site.

### The JM1 segment is responsible for accelerated kinase activation

To investigate which region of the JM domain is responsible for accelerating kinase activation measured for ICD (**Figure 1B & 1C**), we expressed and purified ICD variants in which either JM1 (ICD^ΔJM1^) or JM2 (ICD^ΔJM2^) were deleted (**Figure 3A**). We also generated a construct in which JM2 was replaced with a repeating Gly-Ser linker of equal length (ICD^JM2=GSL^) to account for any steric effects resulting from fusing JM1 directly to the KD (**Figure 3A**). Analysis of the autophosphorylation activity of these constructs revealed that deletion of the JM2 segment had no effect on the initial rates relative to the full ICD, whereas deletion of the JM1 segment abolished the accelerating effect of the JM domain on kinase autophosphorylation activity (**Figure 3B**). The initial activation rate of ICD^ΔJM1^ was equivalent to the KD alone and displayed biphasic activation kinetics (**Figure 3B**). The JM1 segment was also responsible for the accelerated substrate-dependent kinase rate (**Figure 3C**). Curiously, while deletion of JM2 did not alter kinase activity, when this segment was replaced with a flexible GS-linker, autophosphorylation and substrate-dependent rates were slightly elevated (∼1.5-fold) compared to the wild type ICD (**Figure 3B & C**). This suggests that JM1-mediated kinase activation may be regulated by both spatial distance from the kinase and the JM2-encoded sequence.

**Figure 3.**
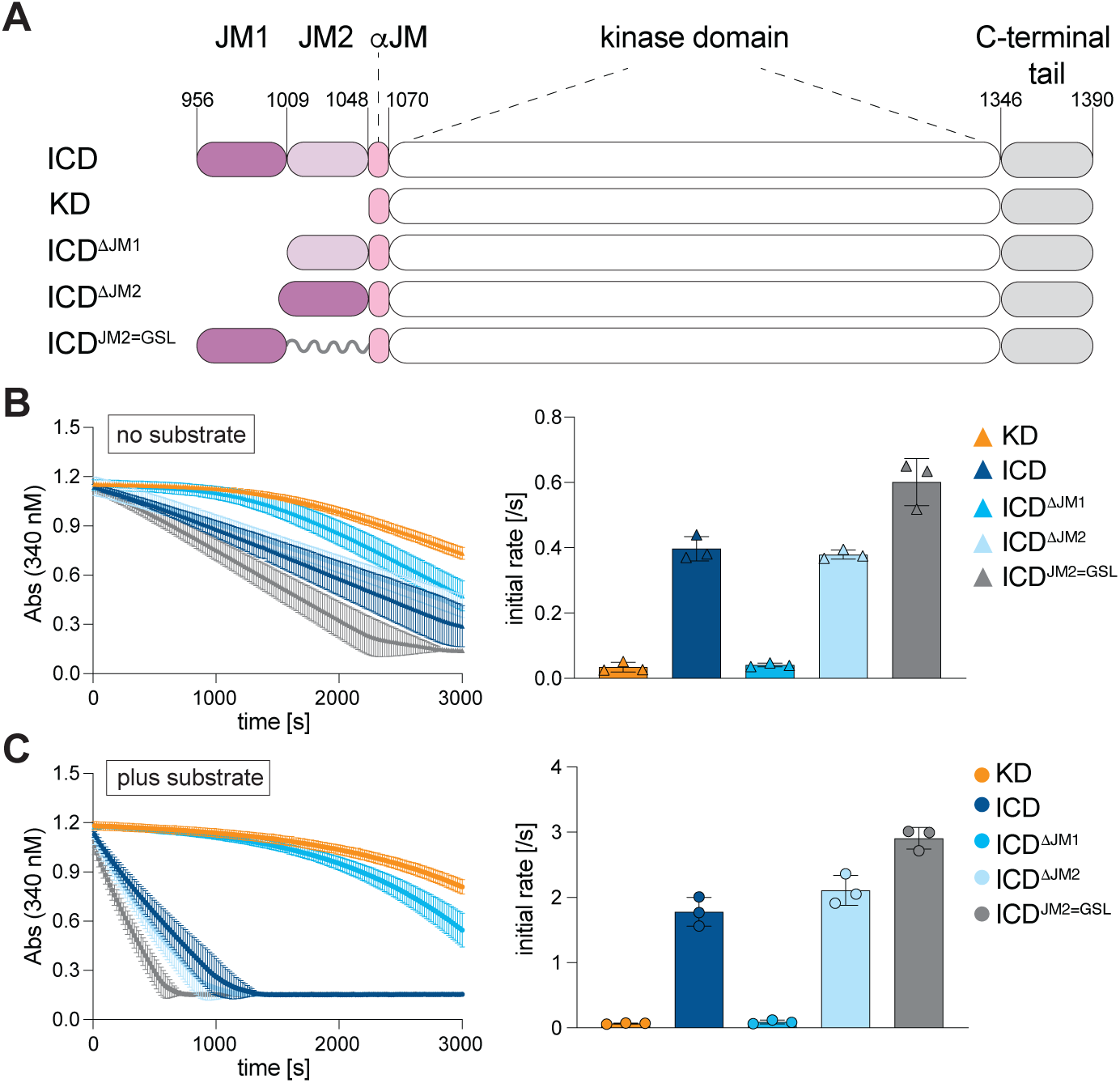
The JM1 region is responsible for accelerated MET kinase activation. (**A**) Schematic depiction of the MET JM domain constructs. (**B-C**) Representative traces from the coupled kinase assay for the dephosphorylated constructs, obtained in the absence (B) or presence of substrate (C). Bar graphs on the right show the calculated catalytic rates from triplicate measurements for a representative experiment. Error bars represent standard deviation (SD). Supplementary Table 1 summarizes average values for each kinase assay from independent experiments.

### Known phosphorylation sites do not directly contribute to the activating effect of the JM1 region

Numerous RTKs are regulated by phosphorylation of tyrosine residues in their JM domains, which frequently serves to release autoinhibitory interactions imposed on the kinase domain (30). We observed that the c-Cbl-binding site Y1003 is autophosphorylated by MET *in vitro* using a phospho-specific antibody (**Supp. Figure 3A**), suggesting that this event may contribute to the accelerated activation in our assays. We investigated whether preventing phosphorylation of this site would impact the potentiating effect of JM1 on kinase activity by generating a non-phosphorylatable Y1003F mutant. We also looked at the effect of mutation of another known JM1 phosphorylation site, the PKC site S985, by generating its non-phosphorylatable variant (S985A) and a phospho-mimetic (S985E). All three mutant ICD constructs exhibited activity equivalent to the wild type ICD (**Supp. Figure 3B - D**), thus arguing against the role of these post-translational modifications in direct regulation of JM1-dependent kinase activation.

### The MET JM domain acts independently of kinase multimerization

One possible mechanism by which JM1 accelerates kinase activation is through promoting multimerization of the intracellular domain, thereby facilitating more efficient trans-phosphorylation of the kinase activation loop. We measured the oligomeric state of the KD and ICD constructs using size exclusion chromatography coupled with multi-angle light scattering (SEC-MALS). The ICD and KD both migrated as single peaks with calculated molecular weights corresponding to their respective monomeric states (**Figure 4A**). Notably, the SEC-MALS analysis was conducted at a protein concentration of 20 µM, 100-fold higher than that in the coupled kinase assay, strongly arguing against JM1-mediated oligomerization as a mechanism of kinase activation.

**Figure 4.**
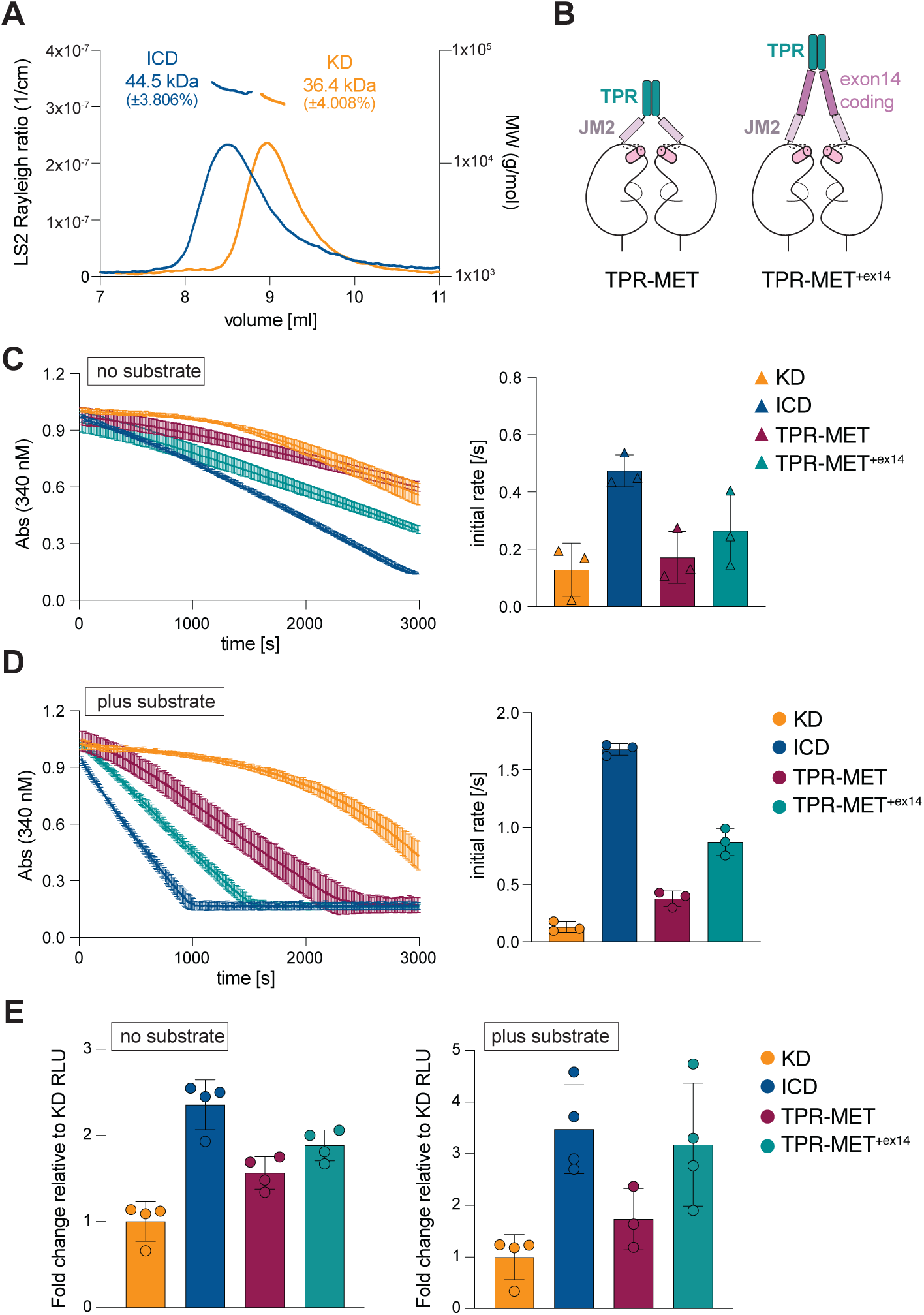
The juxtamembrane domain activates MET independently of dimerization. (**A**) Size-exclusion chromatography coupled with multi-angle light scattering (SEC-MALS) analysis of the dephosphorylated KD and ICD. Elution volume from the size exclusion column is plotted on the x-axis. The left y-axis shows the light-scattering signal. Calculated molecular weights (MW) are indicated above the elution peaks, with their distribution shown on the right y-axis. (**B**) Schematic depiction of the dimeric MET constructs engineered via fusion of the Translocated Promoter Region (TPR). (**C-D**) Representative traces from the coupled kinase assay for the indicated dephosphorylated MET constructs, obtained in the absence (C) or presence of substrate (D). Bar graphs on the right show the calculated catalytic rates from triplicate measurements for a representative experiment. Error bars represent standard deviation (SD). (**E**) ADP-Glo activity measurements for the indicated proteins are shown as fold-change in relative luminescence units (RLU), normalized to the activity of KD measured in the absence (left) or presence (right) of substrate. Supplementary Table 1 summarizes average values for each kinase assay from independent experiments.

Ligand-mediated dimerization is a key step in MET receptor activation, promoting kinase autophosphorylation. Since the isolated ICD is monomeric in our assays, we tested whether the accelerating effect of the JM1 segment is retained when the MET kinase domain is locked in a constitutive dimer. To engineer such dimers, we utilized the gene-fusion product TPR-MET, a rare occurring oncogene, in which a coiled-coil dimerization motif of translocated promoter region protein (TPR) is fused in frame with part of the intracellular region of MET (residues D1010-S1390) (3, 58). This constitutively dimerized MET fragment has been utilized extensively to understand growth factor-independent MET signaling in cells (59–62). Because exon 14-coding region, which encompasses a majority of JM1 segment, is absent in TPR-MET, we generated a variant in which it is restored (residues D963-E1009, denoted as TPR-MET^+exon14^) (**Figure 4B**). All TPR-MET variants were expressed and purified from insect cells followed by lambda phosphatase-mediated dephosphorylation. Surprisingly, the autophosphorylation and substrate-dependent catalytic rate of TPR-MET were only slightly elevated compared to the monomeric KD, indicating that TPR-mediated dimerization of the MET kinase domains does not significantly accelerate their activation (**Figure 4C-D**). The activity of dimerized TPR-MET was also significantly lower than that of monomeric ICD, supporting the conclusion that the JM domain’s potentiating effect on kinase activity is unlikely to stem from enhanced dimerization. Furthermore, the region encoded by exon 14 elevated activity of the dimerized kinase domain (TPR-MET vs TPR-MET^+exon14^) (**Figure 4C-D**), suggesting that the JM1 domain might be activating the kinase via an allosteric mechanism such as direct binding. The lower observed activating effect of the JM in the TPR-MET context, relative to the monomeric ICD, may indicate that TPR-mediated dimer restrains or limits the ability of the JM domain to engage the associated kinase domains. The TPR-MET^+exon14^ construct also lacks the short poly-basic region (residues K956-D963) present in the full MET ICD, thus, it is also possible that this region provides additional activating signal.

We used an ADP-Glo assay as an alternative activity assay to corroborate our measurements obtained using the TPR fusion constructs. Relative kinase activity was determined in the presence of saturating concentrations of ATP/MgCl_2_ and in the presence or absence of the substrate after a 60-minute incubation time. Consistent with the coupled kinase assay results, the ICD had higher activity relative to KD (2.8-fold) (**Figure 4E**). In the ADP-Glo assay, TPR-MET exhibited a measurable (1.8-fold) increase in autophosphorylation compared to the KD (**Figure 4E**), and inclusion of the exon 14-coding region further increased activity (2.5-fold over KD) (**Figure 4E**). Similarly, in the presence of the substrate, exon 14-coding region further accelerated kinase activity of TPR-MET (**Figure 4E**). While differences in the assay format and associated activity readout between the ADP-Glo (endpoint, total ATP hydrolysis) and coupled kinase assay (initial ATP hydrolysis rate) may influence apparent activity differences between the constructs, this analysis collectively highlights that the JM1 segment accelerates MET kinase activation of the monomeric kinase domain, without promoting its dimerization.

### The JM regulatory role is not conserved in RON

MET has only one close homolog, called Recepteur d’Origine Nantais (RON) or macrophage-stimulating protein receptor (MST1R). RON also contains an elongated JM domain (residues 983-1074), although it shares only modest sequence homology with MET (24% amino acid identity) (**Figure 5A**). Previous studies identified two putative regulatory regions within the RON JM domain. The first corresponds to a stretch of 27 amino acids (P1009-V1035), designated as JMB, which inhibits RON activation in cells (63, 64). The second is a short stretch of acidic residues (EDE, E1044-E1046), whose mutation to alanine residues increases RON activation in cells (64). *In silico* analysis of the human RON JM domain using Aminode revealed an ECR corresponding to the JMB segment, while the remainder of the JM domain showed a relatively high substitution score across RON orthologs (**Figure 5B**). AlphaFold3 predicted a short alpha helix in the RON JM, which shows sequence similarity to the equivalent region in MET, indicating a potential overlapping function for this segment (**Figure 5C**).

**Figure 5.**
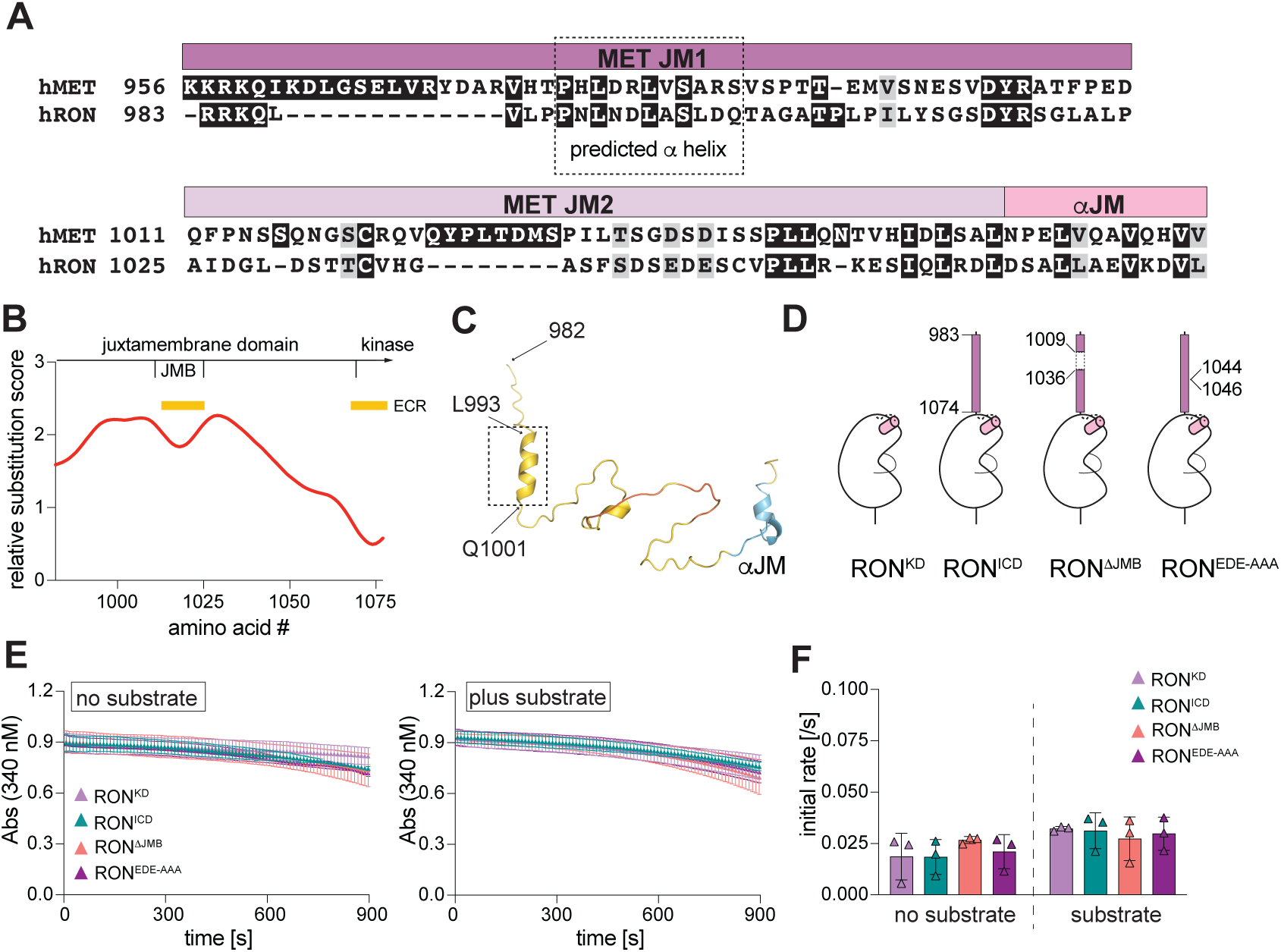
Regulation of RON kinase activity by the juxtamembrane domain. (**A**) Sequence alignment of the human MET and RON juxtamembrane domains with key regulatory regions of the MET JM annotated. The α helix predicted in the N-terminal region of the MET and RON JM domains is outlined. (**B**) Conservation of the RON JM sequence assessed using the *Aminode* algorithm. Values on y-axis represent relative substitution score for each amino acid number, with higher numbers indicating less sequence conservation. Yellow bars mark evolutionary conserved regions (ECRs) within the JM. (**C**) AlphaFold3 model of the RON JM domain, colored by pLDDT (predicted local distance difference test) confidence scores. The location of the predicted α helix is outlined. (**D**) Schematic depiction of generated RON constructs with key amino acids annotated. (**E**) Representative traces from the coupled kinase assay for the indicated dephosphorylated RON constructs, obtained in the absence (left) or presence of substrate (right) (RON kinase was assayed at 1 µM final enzyme concentration). (**F**) Bar graphs on the right show the calculated catalytic rates from triplicate measurements for a representative experiment. Error bars represent standard deviation (SD). Supplementary Table 1 summarizes average values for each kinase assay from independent experiments.

To test if the activating role of the JM domain on MET kinase activity is conserved in RON, we expressed, purified and dephosphorylated RON KD (R1053-T1400) and RON ICD (R983-T1400) fragments (**Figure 5D**). RON KD showed slow biphasic activation kinetics, similar to MET KD (**Figure 5E**). However, RON ICD had equivalent activity to RON KD, indicating that the RON JM domain, unlike MET, does not accelerate kinase autophosphorylation (**Figure 5E & F**). We additionally expressed and purified RON ICD constructs lacking the JMB fragment (RON^ΔJMB^) or with mutation of the acidic patch residues to alanine (RON^EDE-AAA^). In the dephosphorylated form, the activity of these mutants was indistinguishable from the wild type RON ICD, showing that these regions do not directly contribute to catalytic activity of the RON KD. These findings point to a unique mechanism for kinase regulation by the JM domain in MET.

## Discussion

The extended JM domain sets MET apart from most other RTKs, with the exception of its closest homolog, RON, and the unrelated discoidin domain receptor (DDR) RTK family in which JM domains encompass ∼130-160 amino acids (65, 66). Across the RTK family, the JM domain plays essential and versatile roles as a direct modulator of kinase activity.

It can act as an inhibitor, as seen in PDGFR, c-KIT, FLT3, and Eph receptors; as an activator, as in the HER family; or as both, as in the Insulin receptor. Although the MET JM domain contains post-translational modification sites linked to negative regulation of MET receptor signaling by c-Cbl and PKC, to date, no direct role for this domain in controlling kinase activity has been described. Here we report the first direct analysis of the role of the MET JM domain in activation of the kinase domain *in vitro* and show that the N-terminal JM region, denoted by us as JM1, substantially increases initial catalytic rate of the MET kinase by accelerating autophosphorylation kinetics of the activation loop tyrosines Y1234 and Y1235. We found that once the activation loop tyrosines in MET are phosphorylated, the JM domain does not further elevate kinase activity. Thus, the JM domain promotes transition of the MET kinase from an inactive to an active state via accelerating kinase autophosphorylation.

The activating effect of the JM domain maps to its N-terminal 46 amino acids (the JM1 segment), as only the removal of this segment prevented the accelerated activation kinetics in our studies. Our analysis shows that within the JM domain, the JM1 region is most highly conserved. It contains nine serine/threonine and two tyrosine residues known to be phosphorylated, including regulatory sites for c-Cbl recruitment (Y1003) and PKC-mediated downregulation (S985), indicating this region is important for MET receptor regulation. Our computational analysis points to a high likelihood of an α helix in the JM1 region, which encompasses S985. Although the contribution of the predicted α helix to the activating function of the JM1 segment remains undefined, our results demonstrate that mutation of S985 does not alter this activity.

At present, we do not fully understand the mechanism by which the MET JM1 segment accelerates kinase autophosphorylation. In many RTKs, which harbor a short JM domain, extracellular domain dimerization and reorganization of the transmembrane helices helps to bring the kinase domains into close proximity for their transactivation, and the JM domain often regulates this process. This is the case in EGFR, for example, in which JM domain promotes kinase dimerization following growth factor stimulation (19, 36). However, the JM domain in MET presents a challenge for driving kinase proximity, due to its 93 amino acid length, predicted largely unstructured nature and the resulting high conformational flexibility. Consequently, we did not observe a measurable effect of JM domain on MET kinase oligomerization, and the ICD construct remained monomeric far beyond the concentration range used in the kinase assays. Moreover, constitutive MET kinase dimerization in the TPR-MET construct led to only a modest increase in MET catalytic rate, whereas inclusion of the JM1 domain activating segment to this construct elevated kinase activity. Collectively, this suggests that the activating effect of the JM domain might be mediated through a direct interaction with the kinase domain, which could either tether the kinases closer to the membrane or release kinase autoinhibition, thereby accelerating activation kinetics.

In support of the allosteric mechanism come our recent studies in which we conducted deep mutational scan analysis of the MET kinase domain, in a TPR-MET context, and used MET receptor signaling-dependent BaF3 cell survival as a readout (52). This analysis pointed to a direct communication between the exon 14-coding region and the interface between helices αC and αJM within MET kinase domain. Specifically, we observed that mutation of the hydrophobic interface between the αC and αJM helices resulted in loss of function variants, but only when the TPR-MET construct included the exon 14-coding region, hinting to a communication between these distant structural elements. The precise mechanism by which the αC/αJM interface regulates MET signaling or activation has yet to be defined. Rotation of helix αC is a critical step in the activation of many protein kinases, which have evolved diverse mechanisms to regulate it via direct interactions between regulatory elements and the helix αC (19). Whether the JM1 segment engages this interface to regulate the kinetics of MET kinase activation remains to be determined in future studies.

The JM-mediated effect that lowers the activation threshold required for efficient activation loop autophosphorylation and maximal catalytic activity resembles the effect observed with several oncogenic mutations in the MET kinase domain. These mutations, including L1195V, D1228H, Y1230C, Y1235D, M1250T, typically target the intramolecular autoinhibitory interactions within the MET kinase domain (17, 24, 44). Analogous to the effect of the JM domain we have observed here, analysis of these kinase domain mutants *in vitro* showed accelerated autophosphorylation and initial catalytic rates of the MET kinase compared to its wild type counterpart, without an effect on maximal catalytic rate (25). A similar phenomenon has been shown in cells in which the same mutations can lower the activation threshold required for receptor activation but often do not render MET ligand-independent or constitutively active (21–24, 67). This notion of lowering the activation threshold required for kinase activation, rather than resulting in constitutive activity, is also exemplified by recently described mutations in the MET kinase N-lobe identified in patients with hereditary papillary renal cell carcinoma (HPRC) (67); the mutant receptors remain dependent on HGF activation to drive tumorigenesis.

An intriguing question is how the activating effect of the JM1 segment *in vitro* reported here can be reconciled with the pro-oncogenic effect of exon 14 skipping observed in patients, which effectively corresponds to the loss of the JM1 domain in MET. A recent study showed that oncogenic signaling by exon 14 skipping mutant can be recapitulated by Y1003F mutation, which prevents c-Cbl binding within the exon 14 region (47). This suggests that exon 14 skipping exerts its oncogenic signaling primarily by slowing down MET downregulation. This could suggest that slower MET activation kinetics, expected to result from loss of JM1, and prolonged signaling due to loss of c-Cbl binding may cooperate to disrupt intracellular receptor regulation, ultimately promoting cellular transformation. Similar to kinase domain mutations, exon 14-skipped MET remains ligand-dependent in the absence of receptor overexpression, and cells harboring this variant show prolonged signaling upon HGF stimulation (68). An alternative model that rationalizes why exon 14 skipping exerts a net activating effect on MET signaling assumes that shortening the JM domain brings the kinase domains into closer proximity to the plasma membrane, which serves as a hub for interactions of multiple signaling modules, including PI3K and AKT. Receptor dimerization, kinase activation and recruitment of these signaling regulators may coalesce a more potent signaling hub to drive oncogenesis, despite a potential decrease in kinase activation kinetics.

In addition to the significant activating effect of the JM domain on MET kinase activity, our studies also reveal a modest inhibitory role of the MET C-tail. The MET C-tail region encodes two key tyrosine phosphorylation sites (Y1349 and Y1356), forming a so called “bidentate” motif, which is crucial for the recruitment of MET downstream adapters including Grb2, PLCγ, PI3K and Gab1. While the C-terminal tyrosines are essential for MET signaling, previous studies have shown the C-tail may also inhibit MET prior to activation, as the catalytic activity of the MET receptor immunoprecipitated from cells is higher in the absence of the C-tail (51). A crystal structure of the MET kinase domain with a portion of the C-tail (residues G1346-K1360) depicts the tail, including the bidentate motif, as docked at the bottom of the MET kinase C-lobe offering a potential mechanism for intramolecular autoinhibition (13). Analysis of the related RON kinase domain *in vitro* also revealed a direct regulation of its catalytic activity by the C-tail (69). Mutation of the bidentate motif, also present in RON C-tail, to phenylalanines similarly increased RON kinase activity, implicating their role in kinase autoinhibition (69). Thus, MET and RON appear to utilize a similar regulatory mechanism involving their C-terminal tails, which is likely relieved upon autophosphorylation of the key tyrosines in this region. In contrast, we show that RON kinase activity is largely unaffected by the JM domain, as its deletion did not alter activation kinetics of the RON kinase. Notably, the JM and C-tail regions appear to act independently in regulating MET kinase activity, as the JM domain substantially impacts catalytic activity regardless of the presence or absence of the C-tail.

Small-molecule inhibitors of MET have shown clinical efficacy in several human cancers, but resistance mutations often limit their durability (70–74). Elucidating the mechanisms that regulate MET signaling is key to managing resistance and uncovering new therapeutic vulnerabilities. Our finding of the new role of the JM domain in regulation of MET kinase adds to the emerging evidence that the MET JM domain functions as a multiplex platform for integrating both activating and inhibitory signals to control MET signaling. While further studies are needed to elucidate the mechanisms by which the JM domain activates MET kinase, our findings highlight the JM domain as a potential therapeutic target for inhibiting MET signaling. For example, small molecules that block JM1-like interactions with the kinase domain could potentially attenuate activation while preserving c-Cbl–mediated downregulation of MET.

## Experimental Procedures

### Constructs

All MET, TPR-MET and RON kinase constructs used in this study were cloned into a modified pFastBac (Thermo Fisher Scientific) plasmid including an N-terminal 10x Histidine tag and TEV cleavage sequence (SYYHHHHHHHHHHDYDIPTTENLYFQG). All point mutants were generated using QuickChange mutagenesis (Agilent Technologies) following standard protocols. All construct boundaries are listed in Supplementary Table 2.

### Protein expression and purification

All MET and RON kinase domain containing proteins, including TPR-MET variants, were expressed and purified from Sf9 insect cells (Expression Systems) using the protocol detailed below. High titer baculovirus for each construct was generated using standard protocols of the Bac-to-Bac system (Thermo Fisher Scientific) and used to inoculate 1 liter cultures (2×10^6^ cells/mL, 27 °C incubation, 110 rpm shaking) at a 1:40 ratio. Cultures were harvested 48 – 72h later by centrifugation and cell pellets were flash frozen and stored at −80 °C. Pellets were thawed on ice in Lysis Buffer (50 mM HEPES, pH 8.0, 250 mM NaCl, 20 mM imidazole, 5 mM β-mercaptoethanol (BME), 10% glycerol (v/v), with addition of cOmplete™ Mini, EDTA-free Protease Inhibitor Cocktail tablets (Roche) and DNase I (Roche). Resuspended pellets were lysed by homogenization with three passages through an EmulsiFlex-C5 (Avestin) at 10 Kpsi followed by ultracentrifugation at 40,000 x g for 1h. Soluble proteins were filtered through 0.45 µm filter prior to loading to HisTrap HP 1 mL or 5 mL columns (Cytiva). Bound proteins were extensively washed in Nickel Buffer A (20 mM HEPES, pH 8.0, 500 mM NaCl, 5% glycerol (v/v), 5 mM BME and 20 mM Imidazole). Increasing concentrations of Nickel Buffer B (Nickel buffer A with 500 mM Imidazole) up to 15% were used to further remove weakly bound proteins followed by a gradient elution from 15 – 100% Nickel Buffer B. Eluted proteins were monitored by SDS-PAGE followed by Coomassie staining, and fractions containing MET or RON were pooled, diluted 1:1 with Nickel Buffer A, and incubated overnight with TEV protease (1:50 TEV/protein) at 4 °C without mixing. TEV-cleaved protein was re-run on a 1 mL or 5 mL column HisTrap HP, and the flow-through was collected, concentrated, and further purified on a S200 Increase 10/300 GL column (Cytiva) in Size Exclusion Chromatography (SEC) Buffer (20 mM HEPES, pH 8.0, 250 mM NaCl, 2 mM (tris(2-carboxyethyl)phosphine) (TCEP) and 5% glycerol (v/v)). Fractions containing pure MET or RON were pooled, concentrated using 30 kDa MWCO centrifuge filters (Amicon) to 0.5 – 4 mg/mL, flash frozen and stored at −80 °C until assayed. To generate dephosphorylated samples, proteins were incubated with Lambda Protein Phosphatase (NEB) and 10 mM MnCl₂ at 18 °C for 16–20 h. For phosphorylation, proteins were incubated with 2 mM ATP and 10 mM MgCl₂ under the same conditions. After dephosphorylation or phosphorylation, all samples were subjected to SEC, concentrated and stored as described above.

### Coupled kinase assay

Kinase activity was measured using a continuous spectrophotometric assay that couples NADH oxidation to ATP consumption (35, 48). All reactions were performed at 25 °C in the presence of kinase reaction mixture (100 mM HEPES, pH 8.0, 10 mM MgCl_2_, 1 mM PEP, 56 U/mL PK/LDH, 0.3 mg/mL NADH and 2 mM ATP) in the presence or absence of 5 mg/mL poly (Glu:Tyr) 4:1 peptide (Sigma). Reactions were initiated by addition of 0.2 mM recombinant kinase unless stated otherwise in the figure legends. The conversion of NADH to NAD+ was monitored by measuring the decrease in absorbance at 340 nM over time using a Biotek Synergy H1 multimode reader (Agilent). Reaction rates were derived from changes in absorbance over time (35) and calculated kcat s^-1^ values are quoted as mean ± SEM. All kinase assays were performed with technical triplicates with at least two independent experiments using independently prepared protein. A summary of all measurements is provided in Supplementary Table 1.

### ADP-Glo kinase assay

Catalytic activity of MET and TPR-MET constructs was determined using the ADP-Glo kinase assay (Promega). Proteins were diluted to 0.2 µM final concentration in 20 µL reaction volume containing 2 mM ATP and 10 mM MgCl_2_ in the presence or absence of 5 mg/mL final poly (Glu:Tyr) 4:1 peptide. ADP production was monitored after 60 min at 24 °C, following the manufacturer’s recommendations, and luminescence was measured on a Biotek Synergy H1 multimode reader (Agilent). The measured Relative Luminescence (RLU) values are represented as fold-change relative to the RLU values for the KD and are quoted as mean ± SEM (Supplementary Table 1).

### Size Exclusion Chromatography-Multi-Angle Scattering (SEC-MALs)

Dephosphorylated MET proteins (100 µL of 20 µM protein) were separated on a Shodex KW-802.5 column by HPLC with a UV detector (Shimadzu), connected with a miniDAWN MALs detector and an Optilab refractometer (Wyatt Technology) at 4 °C. Molecular weight was determined using the Astra software package (Astra v8.3.0.132 Wyatt Technology).

### Western blot analysis of MET phosphorylation

To monitor MET autophosphorylation, dephosphorylated recombinant proteins were diluted to 1 µM in SEC buffer, and an initial sample was taken (time 0) before adding 2 mM ATP and 10 mM MgCl_2_. Reactions were sampled at the indicated times and diluted 4-fold directly in Laemmli sample reducing buffer to stop the reaction. Samples were separated by SDS-PAGE and gels were either stained with Coomassie for total MET quantification or transferred to nitrocellulose membranes for Western blot analysis. The following phospho-specific antibodies were used to detect phosphorylation of MET Y1349 in the C-terminal tail (#3133, CST) and Y1234/Y1235 in the activation loop (#3077, CST), using enhanced chemiluminescence (Amersham) and a ChemiDoc MP Imaging System (Bio-Rad). Intensity of MET specific phosphorylation levels were calculated relative to total MET protein detected by Coomassie staining and quantified using Image Lab software (v6.1, Bio-Rad). Western blot and Coomassie images from replicate experiments are included in Supplementary Data 1. Phosphorylation at MET Y1003 (CST 3135) was also measured to confirm complete dephosphorylation of the recombinant MET protein (Supp Fig 3).

### *In silico* analysis of evolutionary sequence conservation

Human MET phosphorylation sites cited at least five times in prior publications or data sets were extracted from Phosphosite Plus (55). Sequence conservation and evolutionary constrained regions were derived from Aminode (54) using standard parameters. Data were downloaded and reformatted in Prism (v10.4.1) to focus on the intracellular JM domain of MET and RON. Sequence conservation of MET was additionally assessed by sequence alignment of MET orthologs extracted from https://www.ncbi.nlm.nih.gov/datasets/gene/4233/#orthologs. Sequences were aligned using ClustalOmega (75) and a consensus sequence logo for the MET intracellular region is displayed in Supplementary Figure 2. Sequence alignment from selected species with gaps removed isshown in Figure 2E. Secondary structure prediction and protein folding analysis were performed using Jpred 4 (56) and the AlphaFold3 server (57), respectively, using the intracellular JM regions of human MET (amino acids 956-1071) and human RON (amino acids 983-1074). AlphaFold3 models were analyzed in Pymol (v.3.0.0, Schrodinger). Disorder prediction of the equivalent region of MET was assessed using IUPred2A server (76).

### Data Availability Statement

All data supporting the findings of this study are included in the article and its supplementary materials.

## Supplementary Data

**Supplementary Data 1. Replicate measurements of MET autophosphorylation by Western blot and unedited gels and Western blot images.** Independent replicate experiments showing Western blot analysis of MET autophosphorylation using MET phospho-specific antibodies as detailed in Figure 1D.

**Supplementary Table 1.**
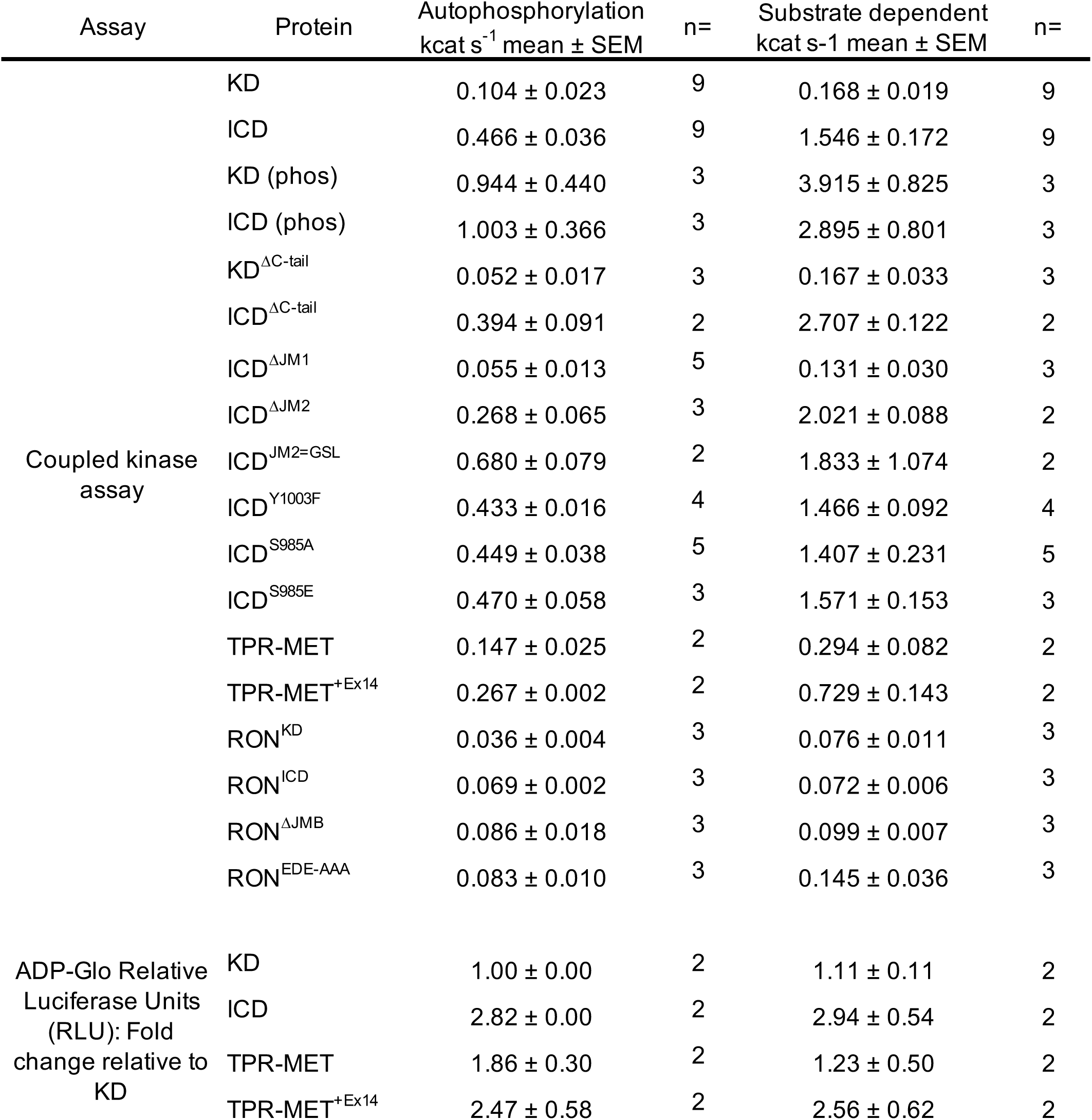
Detailed results of the kinase assays.

**Supplementary Table 2.**
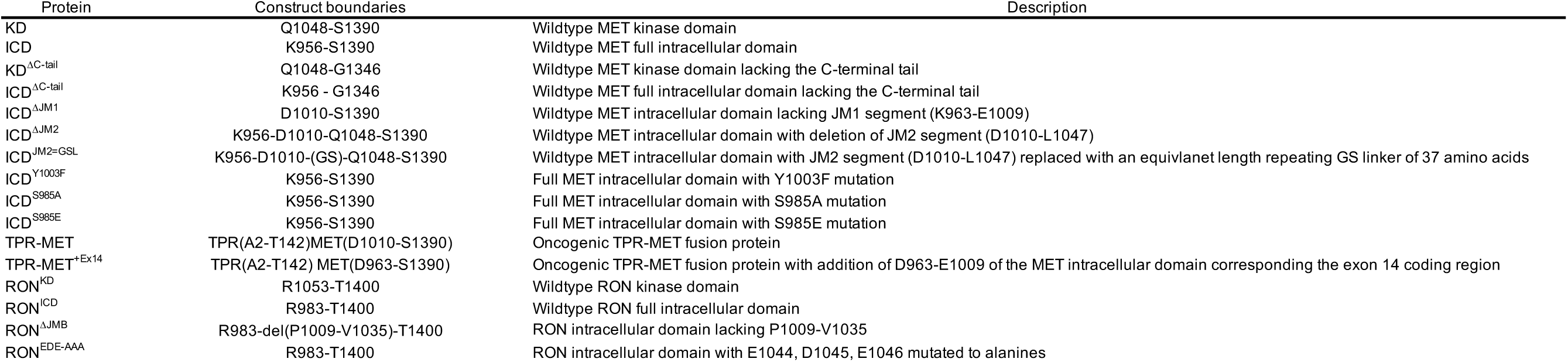
Details of recombinant protein constructs.

## Supplementary Figure Legends

**Supplementary Figure 1. Effect of C-tail deletion on MET kinase catalytic activity.** (**A-B**) Representative data from the coupled kinase assay for the phosphorylated MET constructs, with autophosphorylation rates shown in the absence of substrate (A) and in the presence of substrate (B). Bar graphs on the right show quantified rates from triplicate measurements for a representative experiment. Error bars represent standard deviation (SD). Supplementary Table 1 summarizes average values for each kinase assay from independent experiments.

**Supplementary Figure 2. MET Ortholog consensus sequence logo.** Consensus sequence logo from MET ortholog alignment with MET intracellular domain regions annotated

**Supplementary Figure 3. Mutation of MET JM phosphorylation sites does not alter kinase domain activation.** (**A**) Western blot analysis of the MET ICD before and after incubation with Lambda phosphatase or ATP and MgCl_2_. Phospho-specific antibodies are indicated next to the respective blots. (**B-C**) Representative data from the coupled kinase assay for the indicated dephosphorylated MET constructs, with autophosphorylation rates shown in the absence of substrate (B) and in the presence of substrate (C). (**D**) Bar graph shows quantified catalytic rates from triplicate measurements for a representative experiment. Error bars represent standard deviation (SD). Supplementary Table 1 summarizes average values for each kinase assay from independent experiments.

