## Supplementary Figures and Data for "Autoregulation of the MET Receptor Tyrosine Kinase by its Intracellular Juxtamembrane Domain"

### Supplementary Figure 1

Linossi et al

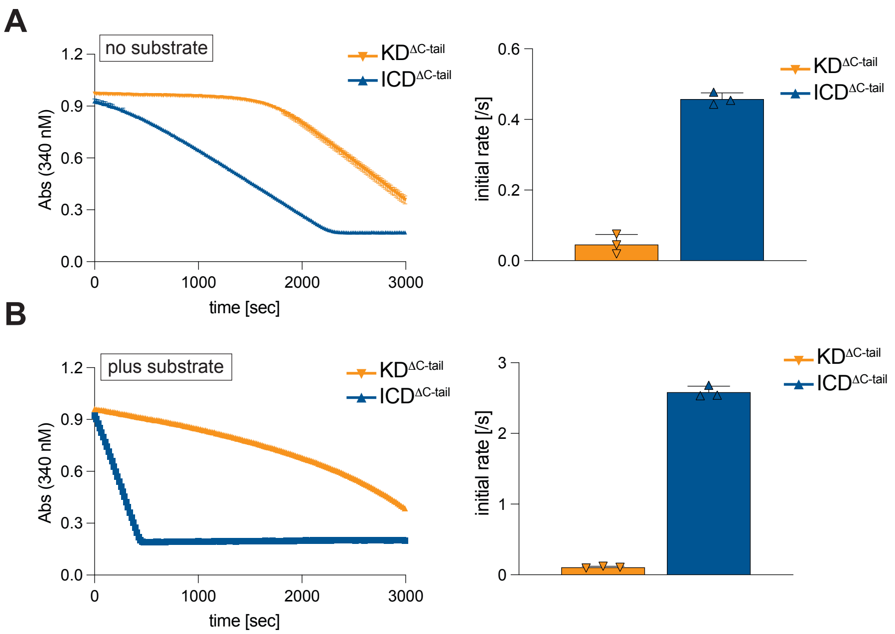

### Supplementary Figure 2

Linossi et al

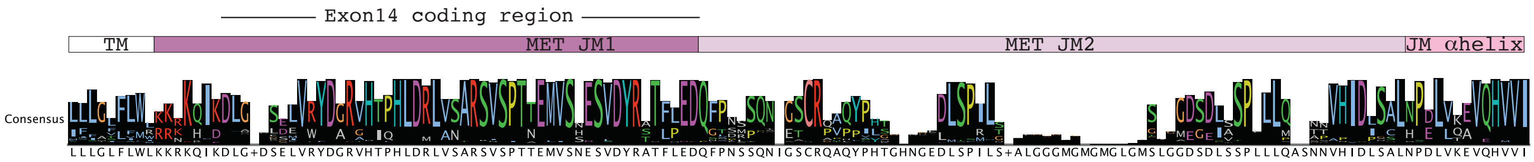

### Supplementary Figure 3

Linossi et al

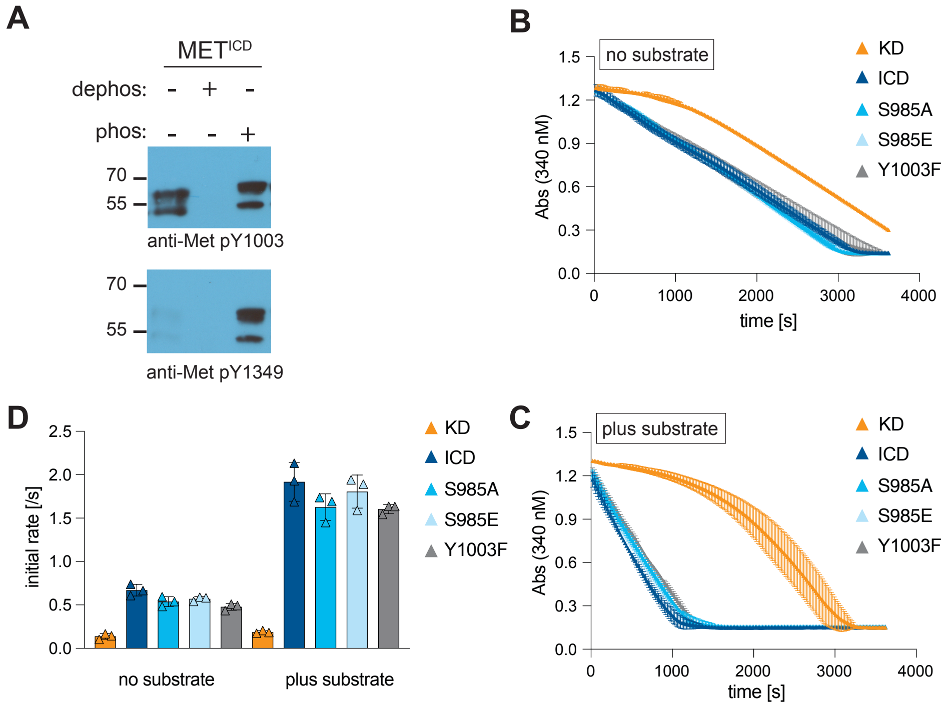

### Supplementary Data 1

Linossi et al

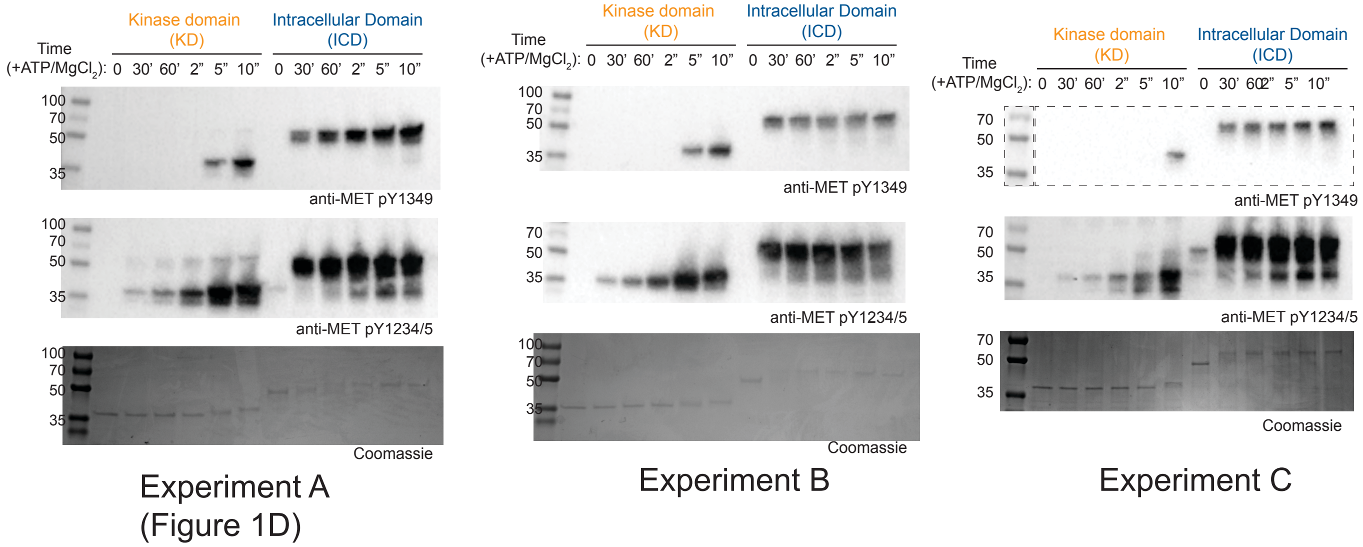


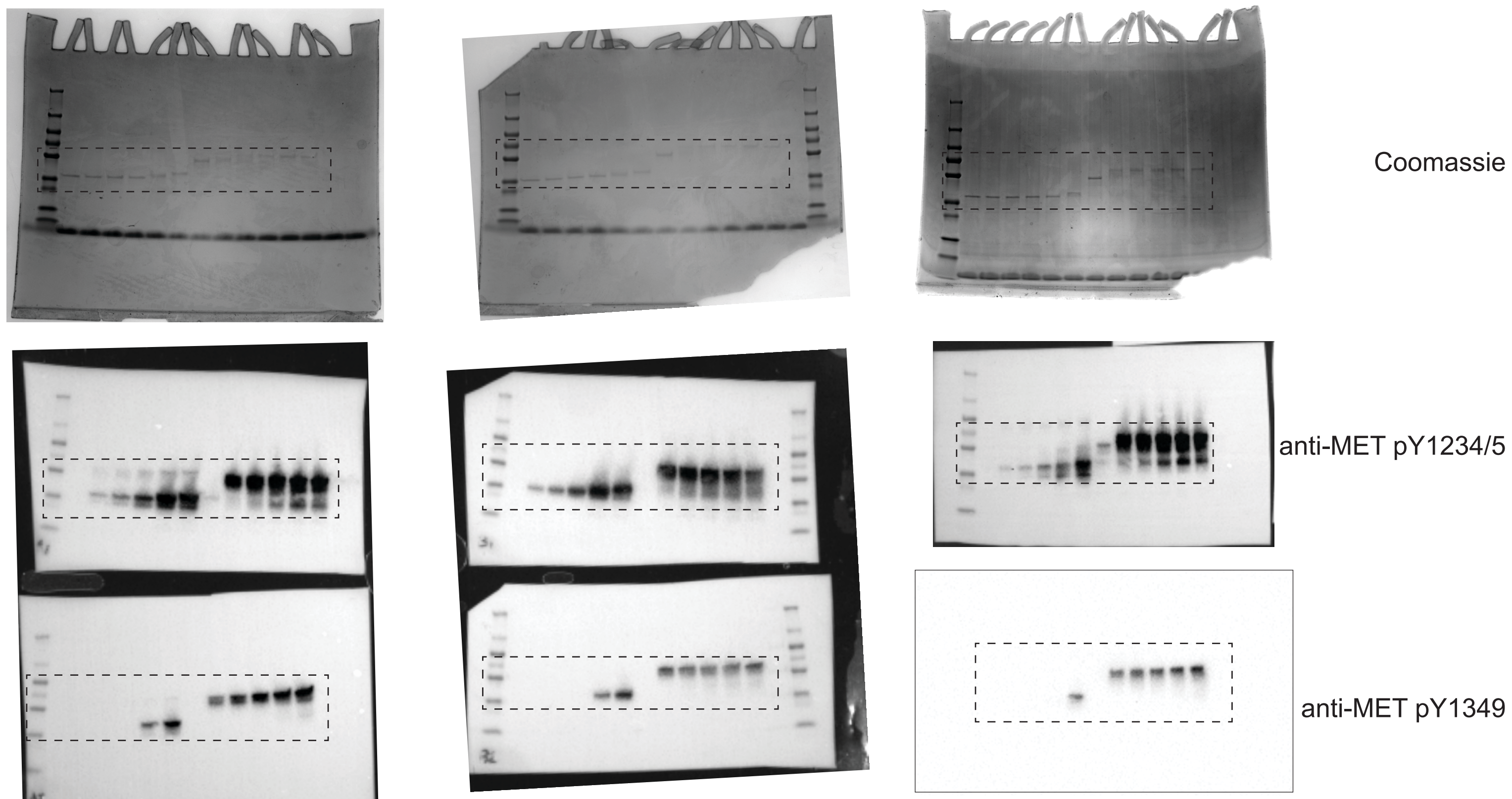


Chemiluminescence image for Experiment C
Boxed area represents region used for Experiment C image
See image below for blot dimensions and edges

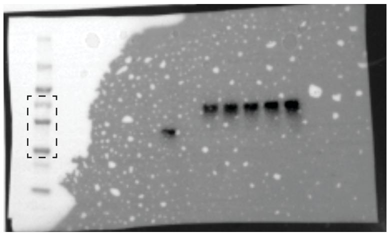

Colorimetric image for molecular weight ladder overlay for Experiment C
Boxed area represents region used for Experiment C image
